# Pervasive mitochondrial tRNA gene loss in the clade B of haplosclerid sponges (Porifera, Demospongiae)

**DOI:** 10.1101/2024.03.04.583380

**Authors:** Dennis V. Lavrov, Thomas L. Turner, Jan Vicente

**Affiliations:** Department of Ecology, Evolution and Organismal Biology, Iowa State University, 241 Bessey Hall, Ames, Iowa 50011, USA; Department of Ecology, Evolution, and Marine Biology, University of California, Santa Barbara, Santa Barbara, CA 93106-9620; Hawai’i Institute of Marine Biology, University of Hawai’i at Manoa, 46-007 Lilipuna Road, Kane’ohe HI 96744-1346

**Keywords:** mitochondrial DNA, tRNA import, comparative genomics, Haplosclerida

## Abstract

Mitochondrial tRNA gene loss and cytosolic tRNA import to mitochondria are two common phenomena in mitochondrial biology, but their importance is often under-appreciated in animals. This is because most bilaterally symmetrical animals (Bilateria) encode a complete set of tRNAs needed for mitochondrial translation. By contrast, studies of mitochondrial genomes in non-bilaterian animals have shown a reduced tRNA gene content in several lineages, necessitating tRNA import. Interestingly, in most of these lineages tRNA gene content appears to be set early in the evolution of the group and conserved thereafter. Here we demonstrate that Clade B of Haplosclerid Sponges (CBHS) represent an exception to this pattern. We determined mt-genome sequences for eight species from this group and analyzed them with six that had been previously available. In addition, we determined mt-genome sequences for two species of haploslerid sponges outside the CBHS and used them with eight previously available sequences as outgroups. We found that tRNA gene content varied widely among CBHS species: from three in an undescribed *Haliclona* species (Haliclona sp. TLT785) to 25 in *Xestospongia muta* and *X. testudinaria*. Furthermore, we found that all CBHS species outside the genus *Xestospongia* lacked *atp9*, while some also lacked *atp8*. Analysis of nuclear sequences from *Niphates digitalis* revealed that both *atp8* and *atp9* had transferred to the nuclear genome, while the absence of mt-tRNA genes represented their genuine loss. Overall, CBHS can be a useful animal system to study mt-tRNA genes loss, mitochondrial import of cytosolic tRNA, and the impact of both of these processes on mitochondrial evolution.

**Significance statement:** It is generally believed that the gene content is stable in animal mitochondrial (mt) DNA. Indeed, mtDNA in most bilaterally symmetrical animals encompasses a conserved set of 37 genes coding for 13 proteins, two rRNAs and 22 tRNAs. By contrast, mtDNA in non-bilaterian animals shows more variation in mt gene content, in particular in the number of tRNA genes. However, most of this variation occurs between major non-bilaterian lineages. Here we demonstrate that a group of demosponges called Clade B of Haplosclerid Sponges (CBHS) represents a fascinating exception to this pattern, with species experiencing recurrent losses of up to 22 mt-tRNA genes. We argue that this group constitutes a promising system to investigate the effects of tRNA gene loss on evolution of mt-genomes as well as mitochondrial tRNA import machinery.

## Introduction

As descendants of the *α*-proteobacterial partner in the primary symbiotic event, mitochondria inherited the eubacterial information processing machinery. While most components of the DNA replication and transcription complexes have been replaced with other, often viral, parts (Shutt and Gray, 2006), mitochondrial translational apparatus has been better preserved (Ott et al., 2016). In particular, nearly all mitochondria contain eubacteria-derived genes for large and small submit ribosomal RNA (*rnl, rns*) and, usually, at least some tRNAs (Salinas-Gieǵe et al., 2015). In addition, multiple nuclear genes of eubacterial origin encoding mitochondrial ribosomal proteins, several mitochondrial amino-acyl tRNA synthethases and two initiation factors are usually retained (Lightowlers et al., 2014; Rudler et al., 2020).

With a few exceptions (Dörner et al., 2001; Barthelemy and Seligmann, 2016), mtDNA in Bilateria (animals with bilateral symmetry) encode a set of tRNAs deemed sufficient for translating all mitochondrial codons (Watanabe and Yokobori, 2011). By contrast, non-bilaterian animals (phyla Cnidaria, Ctenophora, Placozoa, and Porifera) show a large variation in the number of mt-tRNA genes (Pett and Lavrov, 2015). Thus ctenophores lost all mt-tRNA genes (Pett et al., 2011), cnidarians retained at most two (Beagley et al., 1998; Kayal et al., 2012), while studied placozoans species encode a complete set of mt-tRNAs (Miyazawa et al., 2020). Sponges (phylum Porifera) show more diversity in mt-tRNA gene content, which varies from two in the subclass Keratosa (Wang and Lavrov, 2008) to as many as 27 in the homoscleromorph genus *Oscarella* (Wang and Lavrov, 2007). Interestingly, in many animal lineages the number of tRNA genes appears to be set up early in the evolution and preserved thereafter (Lavrov and Pett, 2016). A group of sponges – so-called Clade B of Haploslerida sponges (CBHS) – represents a fascinating exception to the described pattern and displays a large variation in the number of mt-tRNA genes (Lavrov et al., 2023).

Order Haplosclerida Topsent 1928 is a large group of common marine sponges that contains 1,146 described species (de Voogd et al., 2023). Because traditional taxonomy of haploslerid sponges is highly problematic (McCormack et al., 2002), a molecular classification system has been proposed, that subdivides this group into several clades named A-E (Redmond et al., 2011; Redmond et al., 2013). Clade B of Haploslerida sponges (CBHS) is a relatively well-studied group that includes such iconic species as the giant barrel sponge (*Xestospongia muta*), the largest sponge species on Caribbean reefs (McMurray et al., 2008), as well as *Amphimedon queenslandica*, a model sponge species (Srivastava et al., 2010). The mt-genome of *X. muta* is well conserved and encodes a complete set of 25 tRNAs (Wang and Lavrov, 2008). By contrast, mtDNA of *A. queenslandica* encodes only 17 tRNAs, lacks *atp9*, and display faster rates of sequence evolution (Erpenbeck et al., 2007). Our recent study demonstrated that these idiosyncrasies are not restricted to *A. queenslandica*, but appear to be common in the CBHS (Lavrov et al., 2023). In fact, mtDNA in *Niphates digitalis* and *N. erecta* lacks *atp8* in addition to *atp9*, encodes only four tRNAs, and displays even higher rates of sequence evolution (Lavrov et al., 2023).

While previous studies have clearly identified CBHS as a group with unusual mitochondrial evolution, the small number of studied species impeded comparative analysis among them. Here we report ten additional mt-genomes of haplosclerid sponges, eight of them belonging to the CBHS. We investigated patterns of tRNA and protein gene losses as well as their effects on evolution of coding sequences in the group.

## Results

### Genome organization and nucleotide composition

Fourteen complete or nearly complete mt-genomes of CBHS species were analyzed in this study. Six of these genomes have been previously described and eight are new to this study (Table 1). In addition, ten complete mt-genomes from clades A and C were used as outgroups, two of them determined for this study. The latter genomes will be described elsewhere.

**Table 1.** Mitochondrial genomes of CBHS sponges.

| Species name | size | # CDS | # tRNA | %AT | AT-skew | GC-skew |
| --- | --- | --- | --- | --- | --- | --- |
| *Amphimedon queenslandica* | 19,960 | 13 | 17<sup>1</sup> | 56 | -0.02 | 0.28 |
| *Gelliodes wilsoni* | 18,679 | 13 | 24<sup>2</sup> | 65.8 | -0.14 | 0.33 |
| *Haliclona caerulea* | 19,262 | 13 | 16<sup>3</sup> | 55.9 | -0.04 | 0.30 |
| *Haliclona plakophila* | 17,095 | 12 | 5 | 63.7 | -0.15 | 0.30 |
| *Haliclona simulans* | 21,865 | 13 | 23 | 56.5 | -0.08 | 0.25 |
| *Haliclona loe* | 17,075 | 12 | 5 | 63.7 | -0.16 | 0.31 |
| *Haliclona pahua* | 18,595 | 13 | 24<sup>4</sup> | 65.7 | -0.14 | 0.33 |
| *Haliclona* sp. (TLT723) | 18,590 | 13 | 24<sup>4</sup>, 5 | 65.4 | -0.14 | 0.32 |
| *Haliclona* sp. (TLT785) | 16,577 | 12 | 3 | 62.6 | -0.16 | 0.32 |
| *Neopetrosia sigmafera* | 18,188<sup>6</sup> | 13 | 23<sup>4</sup> | 66.1 | -0.15 | 0.34 |
| *Niphates digitalis* | 18,270 | 12 | 4 | 63.4 | -0.09 | 0.41 |
| *Niphates erecta* | 25,173 | 12 | 4 | 63.7 | -0.05 | 0.31 |
| *Xestospongia muta* | 18,990 | 14 | 26<sup>7</sup> | 66.2 | -0.10 | 0.27 |
| *Xestospongia testudinaria* | 18,996 | 14 | 26<sup>7</sup> | 66.1 | -0.10 | 0.27 |

Size, gene content, and nucleotide composition of the CBHS mitochondrial genomes utilized for this study. <sup>1</sup>While 17 tRNA genes had been reported in Erpenbeck et al. (Erpenbeck et al., 2007) *trnN(guu)* and *trnF(gaa)* were not identified with tRNAscan using the Infernal search. <sup>2</sup>*trnS(gcu)* in this species was misidentified by tRNAscan as the second *trnT(ugu)*. <sup>3</sup>*Haliclona caerulea* mt-genomes contains a duplicated *trnS(gcu)* gene. <sup>4</sup>*trnC(gca)*-like sequence was present, but not identified by tRNAscan with Infernal search. <sup>5</sup> *trnL(uag)*-like sequence was present, but not identified by tRNAscan with Infernal search. <sup>6</sup> The region between *cox1* and *nad1* has not been sequenced completely due to the problems with repeats. <sup>7</sup> *trnX(nnn)* was identified in these species, which potentially code for a tRNA-like structure without a complete anticodon arm.

CBHS mt-genomes varied in size between 16,577 and 25,173 bp, had 56-66.2% AT content and encoded 12-14 proteins and 3-25 tRNAs. Among the typical demosponge mt-protein genes, *atp9* was missing in all species outside the genus *Xestospongia*, while *atp8* was missing in the two *Niphates* species as well as *Haliclona plakophila*, *H. loe* and an undescribed *Haliclona* species (TLT785). Among tRNA genes, only *trnMf* and *trnW* were detected by tRNAscan in all species, while *trnMe* was only found in *Xestospongia* spp. (previously *trnI(cau)* was misidentified as *trnM(cau)e* in *A. queenslandica* (Erpenbeck et al., 2007). Mitochondrial gene arrangements in CBHS are shown in Fig 1. As it is generally the case for animal mt-genomes (Lavrov and Lang, 2005), the gene order of major (protein and rRNA) genes was well conserved among the studied species, while both location and presence/absence of tRNA genes were variable among them.

**Figure 1.**
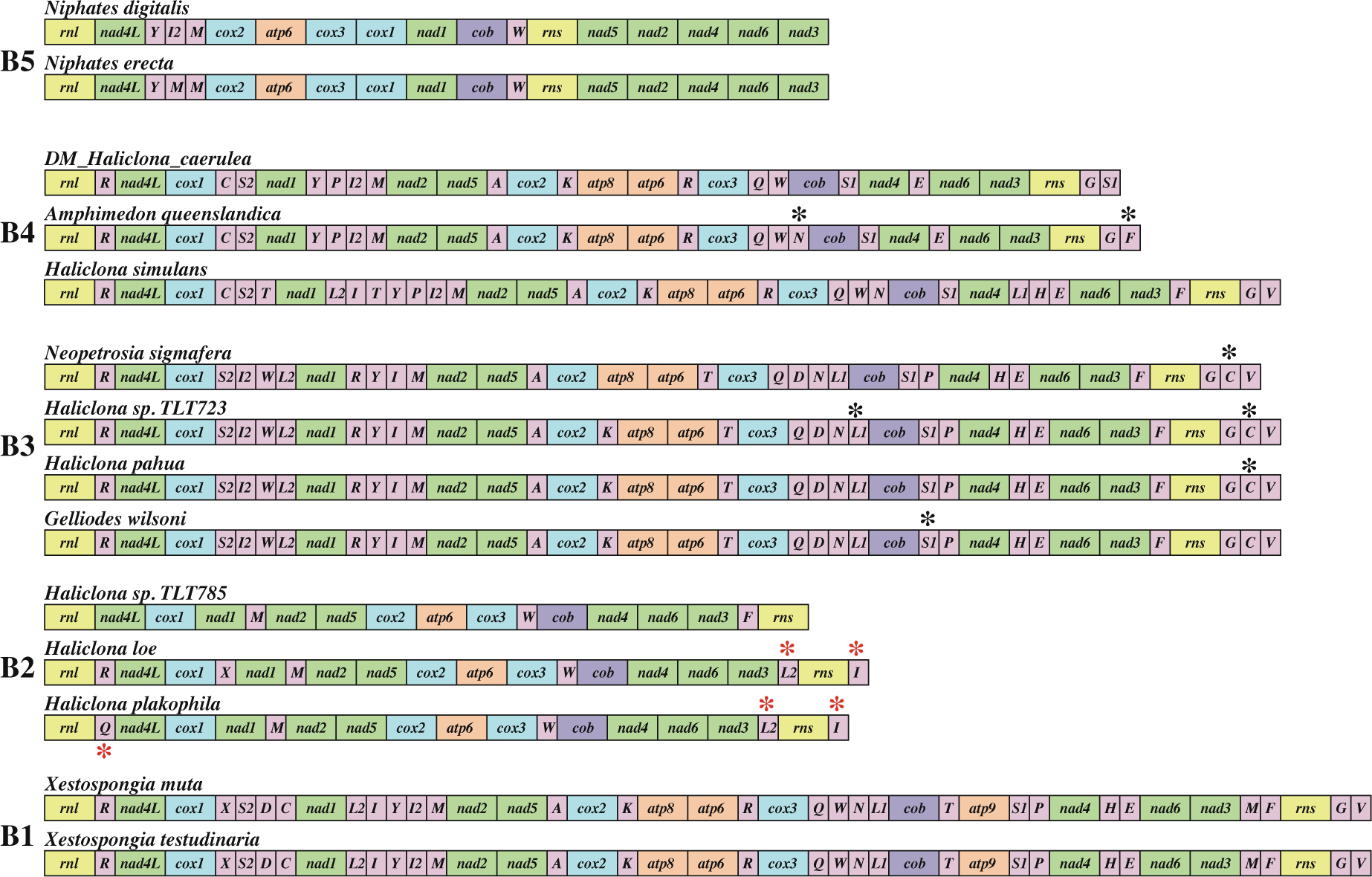
Mitochondrial gene content and gene arrangement in the CBHS. Protein and rRNA genes (larger boxes) are: *atp6*, *8-9* – subunits 6, 8 and 9 of the F0 ATPase, *cox1-3* – cytochrome *c* oxidase subunits 1-3, *cob* – apocytochrome *b*, *nad1-6* and *nad4L* – NADH dehydrogenase subunits 1-6 and 4L, *rns* and *rnl* – small and large subunit rRNAs. tRNA genes (smaller boxes) are abbreviated using the one-letter amino acid code. The two arginine, isoleucine, leucine, and serine tRNA genes are differentiated by numbers with *trnR(ucg)* marked as *R1*, *trnR(ucu)* – as *R2*, *trnI(gau)* – as *I1*, *trnI(cau)* – as *I2*, *trnL(uag)* – as *L1*, *trnL(uaa)* as *L2*, *trnS(ucu)* – as *S1*, and *trnS(uga)* – as *S2*. All genes are transcribed from left to right. Genes are not drawn to scale and intergenic regions are not shown. Red asterisks indicate tRNA genes inferred to evolve by gene remodeling or HGT. Black asterisks indicate genes that were not identified by tRNAscan-SE2 using the Infernal search. They were included in the figure either for consistency with previous studies (*trnN(guu)* and *trnF(gaa)* in *Amphimedon queenslandica* or because of high sequence conservation with closely related species (see below).

The only exception to this pattern were the two *Niphates* species, whose mt-genomes underwent more rearrangements, retaining only four conserved clusters of protein genes: *rnl-nad4L*, *cox2-atp6-cox3*, *cox1-nad1*, and *nad4-nad6-nad3*.

### Phylogenetic analysis of Haplosclerida based on mtDNA data

To track the history of changes in mt-genomes of the CBHS, we reconstructed the phylogeny of Haplosclerida based on concatenated amino acid sequences inferred from mitochondrial protein genes. Our phylogenetic analysis supported the previously proposed clades A, B, and C within the order. Within clade B, five additional clades were recognized, all with the Bayesian posterior probability of 1 (Fig 2):

- Clade B1 consisted of the two *Xestospongia* species and formed the sister group to the rest of the CBHS;
- Clade B2 contained *Haliclona plakophila*, *H. loe*, and an undescribed *Haliclona* species (TLT785);
- Clade B3 encompassed *Neopetrosia sigmafera*, *Gelliodes wilsoni*, *H. pahua* and an undescribed *Haliclona* species (TLT723);
- Clade B4 included *Amphimedon queenslandica*, *H. caerulea*, and *H. simulans*;
- Clade B5 comprised the two *Niphates* species;

**Figure 2.**
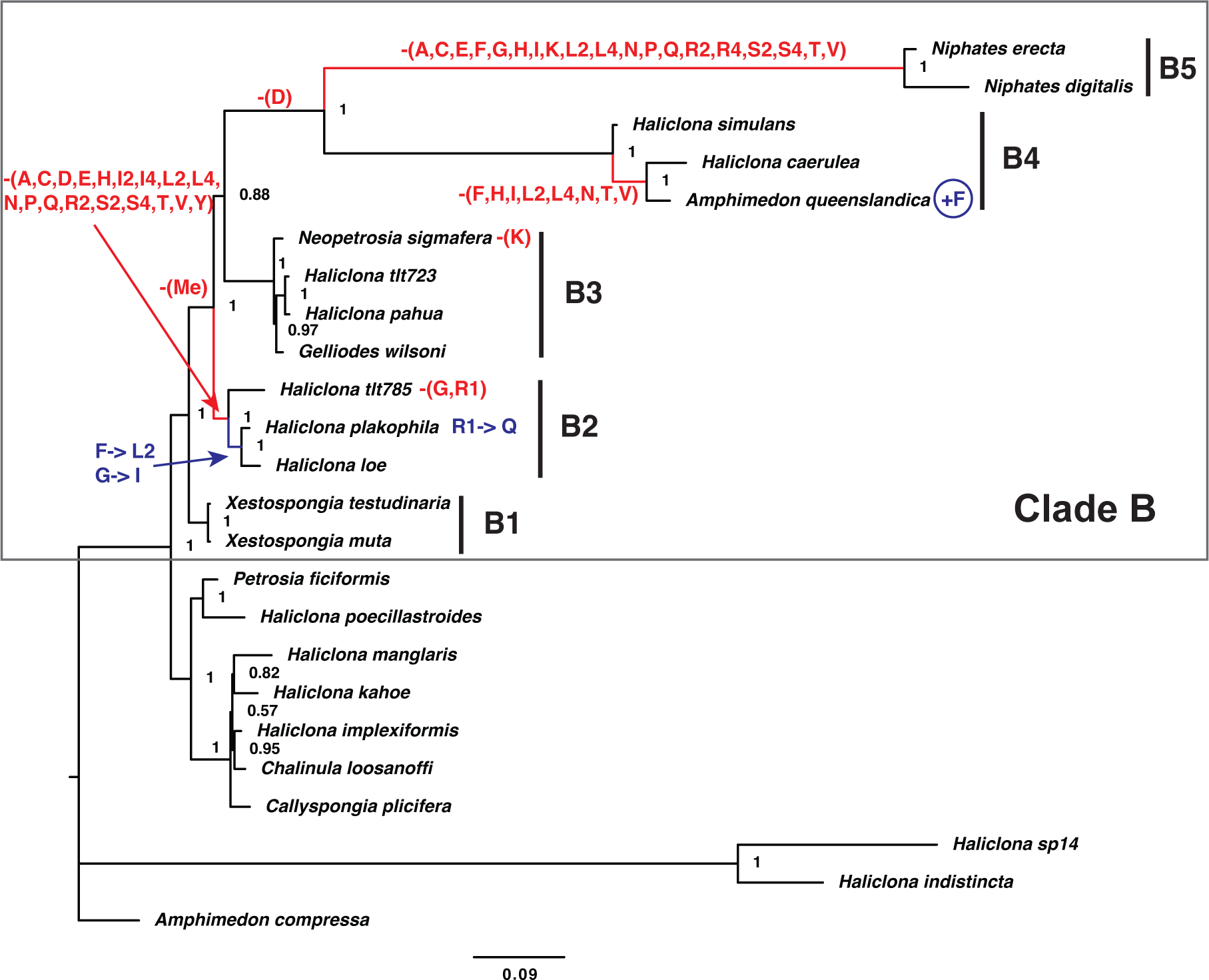
Phylogenetic relationships of haplosclerid sponges with inferred tRNA gene loss and gain. Posterior majority-rule consensus tree obtained from the analysis of concatenated mitochondrial amino acid sequences (3,696 positions) under the CAT+GTR+Γ model with the PhyloBayes-MPI program. The number at each node represents the Bayesian posterior probability. Five major clades in CBHS are shown as B1–B5. We used the Dollo parsimony principle, which assumes the irreversible loss of characters, to manually map gene loss on the phylogenetic tree (in red). The results were adjusted based on the analysis of tRNA gene phylogenetic relationships, which revealed changes in some anticodon identities (in blue). Names of tRNA genes are abbreviated using the one-letter amino acid code.

The inferred phylogenetic tree revealed an accelerated rate of evolution in clades B4 and B5 of the CBHS as well as two species of outgroups.

### Transfer of *atp9* and *atp8* to the nucleus

The most unusual feature of studied CBHS mt-genomes was the absence of several protein and tRNA genes. Mapping observed changes on the phylogeny inferred above suggested that the loss of *atp9* occurred once, in the common ancestor of CBHS species outside the genus *Xestospongia*, while *atp8* has been lost at least two times, in the B2 and the B5 clades (Fig 2). To check whether the absence of mitochondrial genes represents a genuine loss or an intergenomic transfer, we searched for both proteins and tRNA genes in the nuclear genome and transcriptome of *Niphates digitalis*. This analysis identified both *atp8* and *atp9* sequences in the nuclear genome of this species. The encoded ATP8 was 138 amino acids in length and contained a likely mitochondrial targeting sequence (MTS) 41 amino acids in length (TargetP likelihood for MTS 0.86) followed by three additional amino acids prior to the methionine, corresponding to the beginning of this protein in other species. The encoded ATP9 was 141 amino acids in length and contained a MTS of 61 aa in length (TargetP likelihood 0.92), followed by eight amino acids before the first methionine. Because the transfer of *atp9* has been previously reported in *Amphimedon queenslandica* (Erpenbeck et al., 2007), we investigated whether the movement of *atp9* happened in the common ancestor of these species or independently. We found several lines of evidence to support a single translocation of this gene. First, the encoded MTS sequences are 64% identical between the two species, suggesting a single acquisition. Second large sections of each *Niphates digitalis* nuclear contig containing *atp9* was found to be identical to parts of the *A. queenslandica* nuclear contig containing *atp9*. Third, both *atp9* sequences contained an intron in the same position, which was acquired after the transfer to the nucleus. We were also able to identify nuclear contigs containing *atp8* and *atp9* -like sequence in *Haliclona plakophila* DNA-seq data. However because of low coverage of those data, we could not identify them with certainty.

### Loss of mt-tRNA genes and mitochondrial tRNA import

In contrast to protein-coding genes, no mt-tRNA genes were detected in the nuclear genome of *Niphates digitalis*. Thus mt-tRNA genes appear to be genuinely lost. The loss of tRNA genes mapped primarily to three branches: 19 tRNA genes were lost in the B5 clade, 18 tRNA genes were lost in the B2 clade, and eight tRNA genes were lost on the branch leading to *Amphimedon queenslandica* and *Haliclona caerulea* in the B4 clade (Fig 2). A few additional tRNA genes have also been lost in individual species of sponges. Because the loss of a tRNA gene should be preceded by an import of corresponding cytosolic tRNA, one would expect some mt-tRNAs to co-exist and co-function with imported cy-tRNAs. The import of cytosolic tRNAs should lead to relaxed selection on mt-tRNA genes, accumulation of mutations, and eventual disappearance of these genes from mt-genomes. One example of this process was observed with *trnN(guu)* in the clade B4. tRNA encoded by this gene is highly conserved in *H. simulans* (Infernal score 62.6), but contains several mismatches in *A. queenslandica* and is only recognized by tRNAscan with the Cove search option (Cove score 24.5). The same gene accumulated even more mutations in *H. caerulea*, the sister lineage of *A. queenslandica* and was only recognizable by sequence similarity but was not found with tRNAscan (Fig 3). To check for such relaxation of selection on other mtDNA-encoded tRNAs, we compared the tRNA covariance scores calculated by tRNAscan-SE2 in species with and without mt-tRNA loss/cy-tRNA import. We found that the average scores were lower in all species with mt-tRNA loss as compared to *Xestospongia* species, which encode a full set of mt-tRNAs. Similarly, for all tRNAs but *trnA*, the average covariance scores were lower than those in *Xestospongia* species (Fig 4).

**Figure 3.**
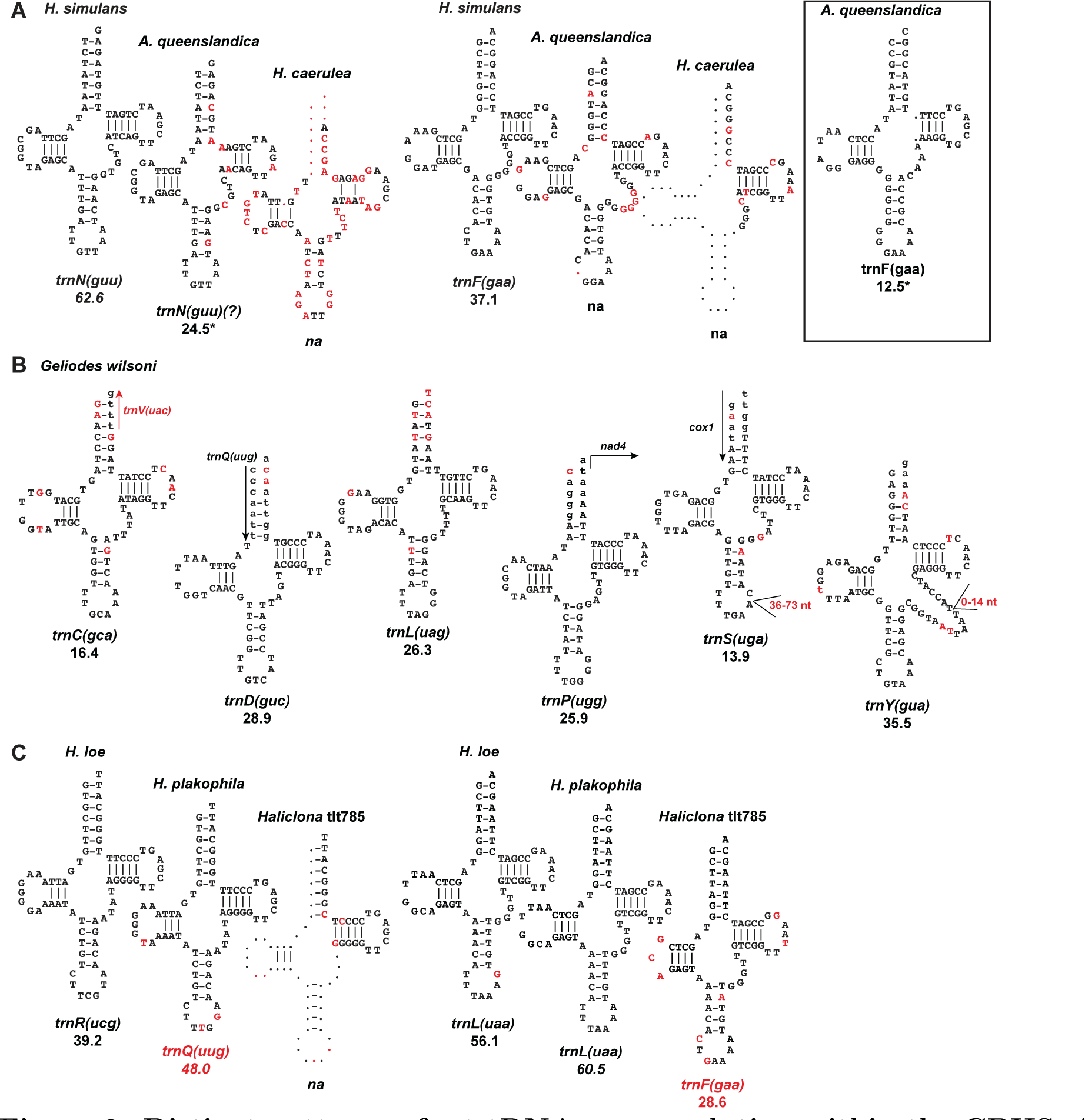
Distinct patterns of mt-tRNA gene evolution within the CBHS. A: Accumulation of mutations leading to losses of mt-tRNA genes within the B4 clade. Nucleotides in red differ from those in *Haliclona simulans*. B: Conservation of unusual tRNA structures within the B3 clade. tRNA sequences are shown for *Gelliodes wilsoni* . Nucleotides in red showed variation among B3 species. Nucleotides in lowercase were not inferred to be part of tRNA by tRNAscan and are shown here as would be expected under the normal cloverleaf structure. C: tRNA remolding in the B2 clade. Remolded tRNA names are in red. Nucleotides in red show differences from corresponding positions in *Haliclona loe*. Number under tRNA names signify covariance score, except when marked with an asterisk. Numbers marked with asterisks signify COVE scores.

**Figure 4.**
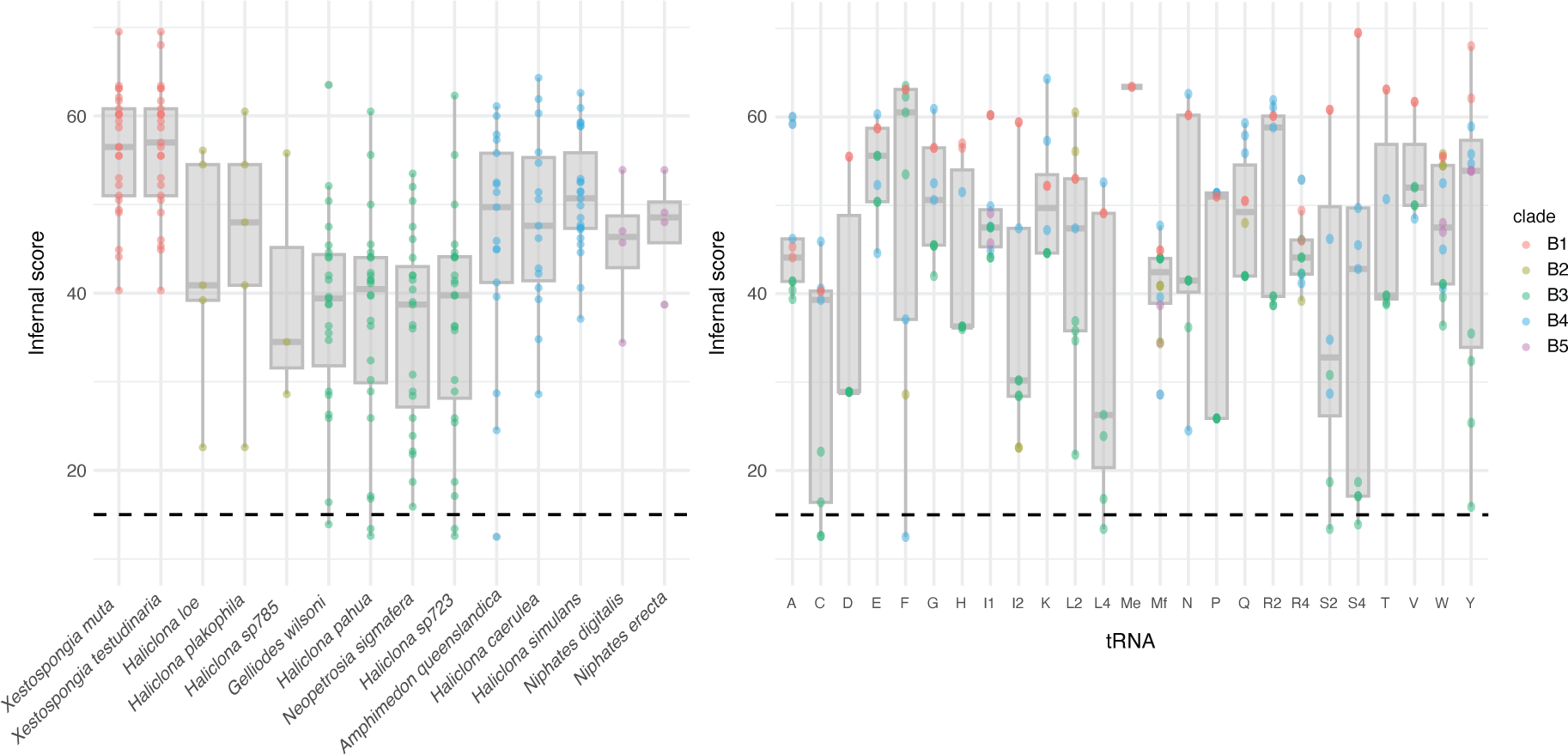
Distribution of tRNA scores in CBHS. Covariance model bit scores for inferred mt-tRNAs in CBHS species are drawn arranged by species (A) and by tRNA The color of the points denotes species’ placement within individual clades of CBHS. The box is drawn from the 25th to the 75th percentile with a horizontal line denoting the median.

### Variation in mt-tRNA gene content, tRNA structural conservation among the subclades of clade B

While we have mainly treated CBHS as a single unit, there was a noticeable variation in the patterns of tRNA loss and evolution among the five identified clades within this group. In particular, haplosclerid species in the clades B2 and B5 lost most tRNA genes, while those in clade B3 retained all but a few. Clade B4 showed more variation in the number of mt-tRNA genes, with *Haliclona simulans* retaining a nearly complete set, while *Amphimedon queenslandica* and *H. caerulea* loosing about a third of them.

Interestingly, there was also a variation in the tRNA structural scores among the clades, with those in the B3 clade showing lower scores for many tRNAs. Several tRNA genes within B3 clade either encoded multiple mismatches in the amino-acyl acceptor stems and/or overlapped with adjacent genes on either 5’ or 3’ end (Fig 3). These included *trnD(guc)*, *trnC(gca)*, *trnL(uag)*, *trnP(ugg)*, *trnS(uga)*, *trnY(gua)*, all with low covarance scores (13.9–35.5). Despite these unconventional structures and low structural scores, tRNA sequences were usually well conserved among the four species within this group. Surprisingly, the best conserved sequences (100% identity among the four species) were those of tRNAs with the most derived structures, such as *trnD(guc)* and *trnP(ugg)* (Fig 3).

### Other changes in mt-tRNA gene family

So far, we considered evolution of tRNA gene family as a function of tRNA gene loss. However, tRNA gene gain, as the result of either horizontal gene transfer or gene recruitment/remolding can contribute to the observed variation in gene content. To check for contribution of these additional factors to tRNA gene family evolution, we constructed a NJ tree of all mt-tRNA sequences identified in CBHS species (Fig 5). Consistent with our previous studies (Lavrov and Lang, 2005; Wang and Lavrov, 2011), we found that most tRNAs for the same codon family clustered together with strong support, while interrelationships among these clusters were only weakly supported. We also found several exceptions to this general pattern:

– First, *Haliclona plakophila* and *H. loe trnL(uaa)* grouped within the clade of *trnF(gaa)* genes from other species and were located in the same genomic position as *trnF(gaa)* genes. No *trnF(gaa)* gene was identified in either of these two species.
– Second, *trnI(aau)* from *H. plakophila* and *H. loe* clustered with *trnG(ucc)* genes from other species with moderate support and was also located in the same genomic position.
– Third, *Gelliodes wilsoni trnS(gcu)* did not group with corresponding genes in other species. This gene was misidentified by tRNAscan as the second *trnT(ugu)* and, hence, was likely misaligned by the cmalign program, which used the same covarance model as tRNAscan.
– Fourth, *H. plakophila* had an unusual *trnQ(uug)* gene that appeared to evolve from the *trnR(ucg)* gene and was located in the same position.
– Fifth, the two *Xestospongia* mt-genomes encoded unusual tRNA-like structure, without an anticodon loop.
– Sixth, there were multiple gene recruitments between *trnT(ugu)* and *trnR(ucu)*.

**Figure 5.**
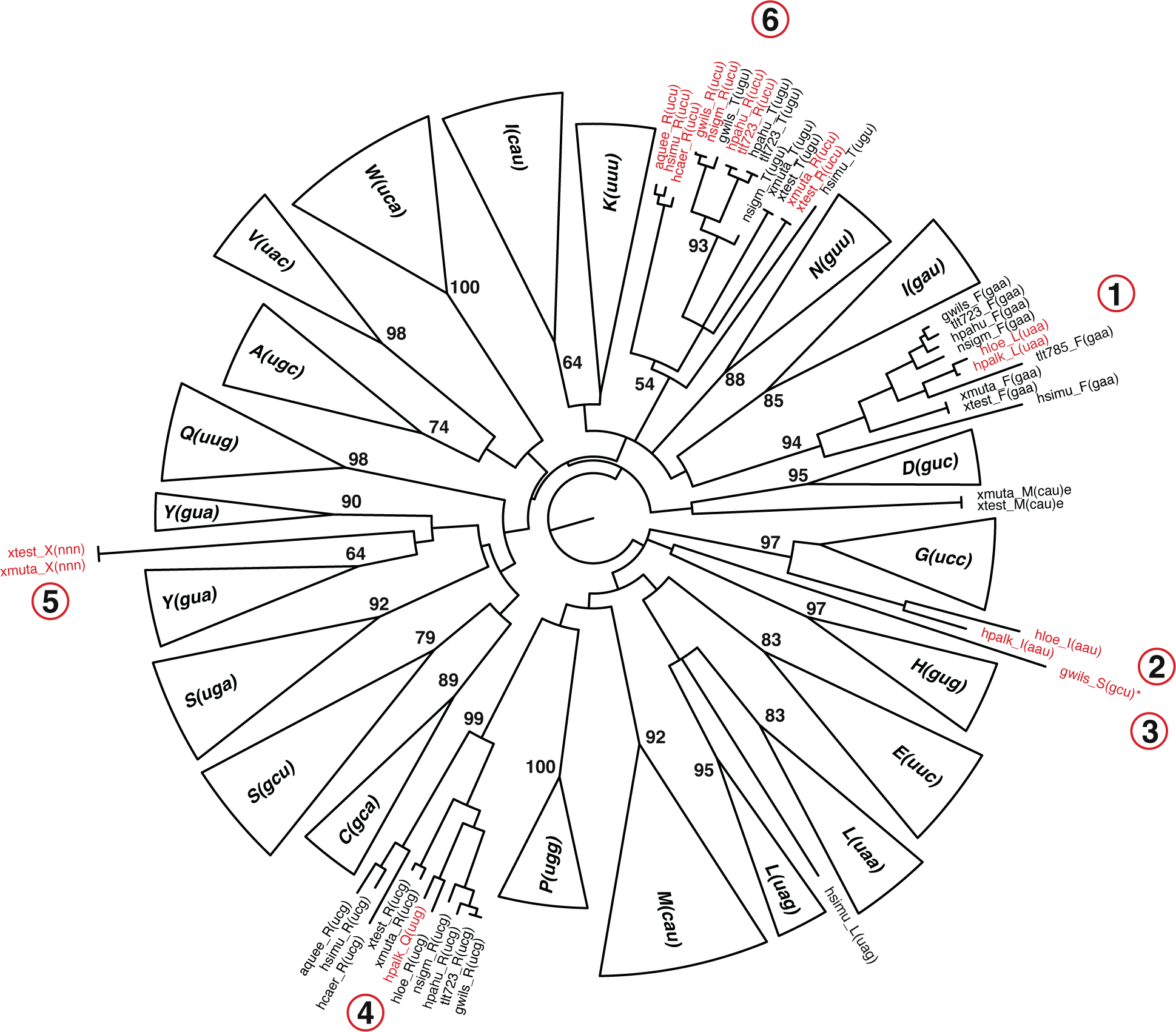
Phylogenetic tree of mt-tRNA genes within the CBHS. Neighbor-joining tree based on uncorrected (P) distances is shown. Clusters containing all and only tRNAs for the same codon family were cartooned. When tRNAs from individual species are shown, species names are abbreviated as following: aquee=*Amphimedon queenslandica*, gwils=*Gelliodes wilsoni*, hcaer = *Haliclona caerulea*, hloe=*H. loe*, hpahu=*H. pahua*, hplak=*H. plakophila*, hsimu=*H. simulans*, nsigm=*Neopetrosia sigmafera*, tlt723=*Haliclona* sp. (TLT723), tlt785=*Haliclona* sp. (TLT785), xmuta=*Xestospongia muta*, xtest=*X. testudinaria*. Numbers above selected branches indicate bootstrap support percentage based on 1000 bootstrap replicates. Circle numbers correspond to listed cases of unexpected phylogenetic results described in the text.

Several of the observations above indicate potential gene gains. Noticeably, all changes in tRNA anticodon identities listed above occurred within the B2 clade: *trnF(gaa) → trnL(uaa)*, *trnG(ucc) → trnI(aau)*, *trnR(ucg) → trnQ(uug)*. However, unlike previous examples of this process in sponges (Lavrov and Lang, 2005; Wang and Lavrov, 2011) as well as the recurrent gene recruitment between *trnT(ugu)* and *trnR(ucu)* observed here, the genes had changed their identities in place, without prior gene duplication. *trnF(gaa)* in *A. queenslandica* identified by Erpenbeck et al. (Erpenbeck et al., 2007) appears to represent another case of tRNA gene remolding as it grouped with *trnG(ucc)* from other species (not shown) and shares identical mt-genome position with them. Furthermore, a degraded copy of the original *trnF* gene was found in its ancestral location (Fig 3).

However, this gene was not identified by tRNAscan using the covariance model and was not included in the phylogenetic analysis shown in Fig 5.

### Did tRNA loss influence the evolution of mt-coding sequences?

Because most animal mitochondrial genomes encode a minimal possible number of tRNAs for a given genetic code, loss of mt-tRNA genes should be preceded and compensated by the import of corresponding cy-tRNAs. However, it is currently unknown what iso-acceptor cy-tRNAs are imported to mitochondria and whether their concentration matches those of mtDNA-encoded tRNAs. Thus, it is possible that the loss of mt-tRNAs would lead to changes in amino-acid usage in the proteins or the changes in codon usage for some amino acids. Pairwise analysis of amino acid composition using Composition profiler (Vacic et al., 2007) showed few changes in amino acid composition in CBHS species with tRNA loss in comparison to *Xestospongia muta*. In particular, no significant changes were observed in the species in the B2 clade, which lost most mt-tRNA genes. However, only a few significant changes were observed in the B4 and B5 clades: the frequencies of Ile and Phe were decreased, while the frequencies of Met, Gly and Val were increased. Those were more likely associated with smaller

AT-skew observed in both B4 and B5 and lower %AT observed in B4 than mt-tRNA gene loss. Consistently, loss of several tRNA genes on the branch leading to *Amphimedon queenslandica* and *Haliclona caerulea* within the B4 clade did not lead to significant changes in amino acid composition of mitochondrial proteins in these species in comparison to *H. simulans*. Similarly, most of the changes in synonymous codon usage were observed between the B4 clade and the rest of the CBHS, with the main difference being higher proportion of cytosine ending codons in the former group (Fig 6).

**Figure 6.**
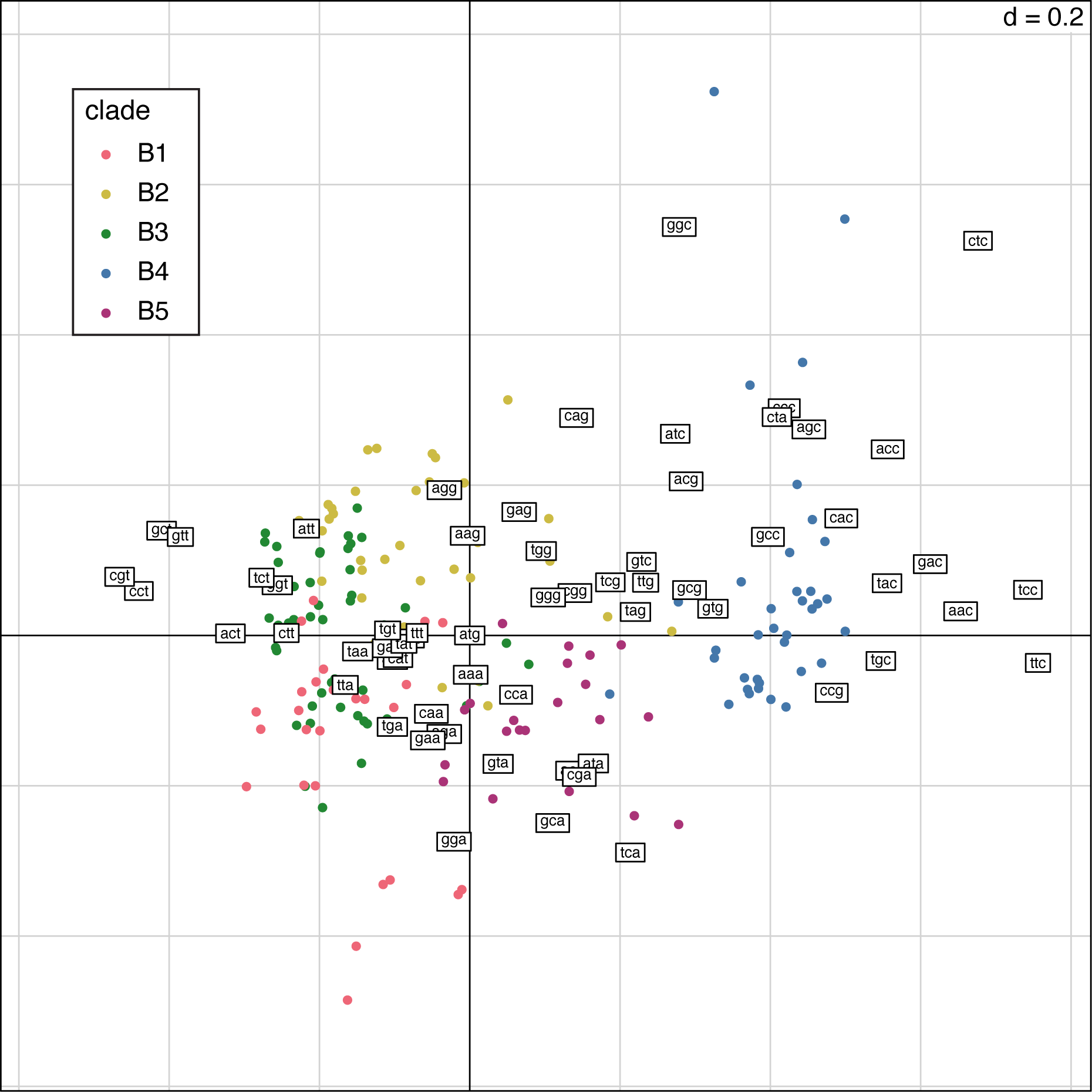
First factorial map for synonymous codon usage in CBHS. Within-group correspondence analysis was conducted for species in Clade B. Each point corresponds to a coding sequence. Species are colored based on their subclade.

Interestingly, T *→* C mutations were also the most common changes in tRNA structures within this group (Fig 3). Thus it appears that the evolution of protein sequences in CBHS is driven more by changes in mutational biases than loss of mt-tRNAs and import of cy-tRNA.

## Discussion

### CBHS as a model to study mt-tRNA gene loss and mitochondrial tRNA import in animals

In contrast to protein import to mitochondria – an extensively studied process with several well characterized pathways – tRNA and, more broadly, nucleic acids import to mitochondria is underexplored (Schneider, 2011). In part, this is due to the fact that mt-tRNA gene loss is not ubiquitous across genes and species and hence tRNA import likely evolved independently in multiple lineages (Schneider and Maŕechal-Drouard, 2000). While in some of them, like yeast (Tarassov and Entelis, 1992) or kinetoplastid protozoa (*Leishmania* and trypanosomes) (Mahapatra and Adhya, 1996; Nabholz et al., 1999; Bhattacharyya et al., 2003) tRNA import mechanisms are relatively well characterized, this knowledge is not directly transferrable to other species (Rubio et al., 2008). At the same time, there is a substantial interest in the process of tRNA import in humans because it has been suggested as a possible avenue for treating mitochondrial diseases caused by mutations in mt-tRNA genes (Kolesnikova et al., 2004) and because the machinery of tRNA import can be recruited to import other nucleic acids to mitochondria allowing genetic manipulation (Wang et al., 2012). Because sponges are phylogenetically more closely related to humans than most other systems where tRNA import has been studied, it is more likely that at least some parts of the machinery involved in mitochondrial tRNA import are conserved between them. While mt-tRNA gene loss and, by inference, mitochondrial tRNA import is present in several groups of sponges (Pett and Lavrov, 2015), CBHS is the only known group that displays a large variation in the tRNA gene content among species. Here we demonstrated that the number of mt-tRNA genes in CBHS species varies between three and 25 and that tRNA loss has occurred repeatedly in this group. This variation can be utilized in comparative studies for characterization of tRNA import machinery in mitochondria. Thus, this group represents a promising model to study mt-tRNA gene loss in animals as well as import of cy-tRNA to mitochondria.

### What constitutes a mt-tRNA gene loss?

Our inference of a tRNA gene presence was primarily based on its detection by the tRNAscan program. While this is a common approach in the analysis of tRNA gene families, it has some caveats. A functional tRNA might have a derived structure and, hence, a low covariance score as commonly seen in mitochondria of bilaterian animals. In contrast, a tRNA with a critical mutation may be otherwise well conserved and have a high score. The ambiguity with tRNA gene annotation can be illustrated by *Amphimedon queenslandica trnN(guu)*. This gene was annotated in the original publication of *A. queenslandica* mt-genome (Erpenbeck et al., 2007) but was not detected by tRNAscan search using covariance model in our study. While the ultimate proof of tRNA functionality would be a demonstration of its participation in protein synthesis, we note that this gene has multiple substitutions in comparison to its ortholog in *Haliclona simulans*, resulting in multiple mismatches in encoded tRNA arms and suggesting a relaxed selection on its function. One example of a low-scoring tRNA that is highly conserved is *trnD(guc)* in the B3 clade, which has a low Infernal score, but identical sequences among the four species in the clade. Thus, supplementing tRNAscan search with comparative analysis may provide a more robust approach to evaluating tRNA gene functionality. It should be noted that considerations above are mostly focused on the primary function of tRNA genes in protein synthesis. However, tRNAs may have additional roles in mitochondrial biology, including their role in mitochondrial ribosomes as well as their participation in mRNA processing (Greber et al., 2014). Thus, a conservation of a tRNA gene sequence may be due to selection on those secondary functions. Another complication in inferring tRNA gene loss and retention comes from tRNA genes that appear to have changed their amino-acid identities. While this process is relatively common in sponges (especially common between *trnT(ugu)* and *trnR(ucu)*, where tRNA amino-acyl identity depends primarily on the anticodon sequence) (Wang and Lavrov, 2011), whether all or any such ”remolded” tRNAs are functional in protein synthesis in CBHS remains unknown.

### Is there a correlation between tRNA loss and other changes in mitochondrial genomes?

In several cases, the loss of mt-tRNA genes in animals is associated with higher rates of mt-sequence evolution. For example, ctenophores lost all mt-tRNA genes and also have one of the highest rates of mt-sequence evolution in animals (Pett et al., 2011). Among demosponges, some of the highest rates of sequence evolution are found in representatives of the subclass Keratosa, which lost all but two mitochondrial tRNAs (Wang and Lavrov, 2008; Erpenbeck et al., 2009). Among homoscleromorph sponges family Plakinidae has higher rate of sequence evolution than Oscarellida and also encode an incomplete set of tRNAs (Gazave et al., 2010). Among haplosclerid sponges, *Niphates* species have some of the highest rates of sequence evolution and also lost most tRNA genes. However, the pattern is not universal. Even among haplosclerid sponges, *Haliclona* sp. 11 has lost as many tRNAs as *Niphates* but does not show accelerated rate of sequence evolution (Fig 2). All representative of the phylum Cnidaria encode at most two mitochondrial tRNAs, but Anthozoa have some of the lowest rates of sequence evolution among animals (Shearer et al., 2002). Thus, while many fast-evolving mt-genomes have lost tRNA genes, not all mt-genomes that lost tRNA genes have high rates of sequence evolution.

Another potential correlation is between losses of mt-tRNA and mt-protein genes. The presence of *atp9* is one of the distinguishing features of sponge mt-genomes, as it is found in all four classes of sponges, but not other Metazoa, where it is located in the nuclear genome (Lavrov, 2007). Nevertheless, there have been several reported losses of this gene from mt-genomes in Demospongiae, including within the CBHS (Erpenbeck et al., 2007; Lavrov et al., 2023), which suggest intergenomic transfers. The transfer of *atp9* to the nuclear genome is interesting, because the protein encoded by this gene forms an oligomeric c ring that make up the Fo rotor of the ATP synthase, and thus requires more copies than other subunits encoded by *atp6* and *atp8* (Stock et al., 1999). Thus it is possible that the transfer of *atp9* to the nuclear genome simplifies expression of mt-genome. In contrast to *atp9*, *atp8* is found throughout Metazoa, but has been independently lost in many species, including some molluscs, arrow worms, nematodes, ctenophores, placozoans, and calcareous sponges (Pett and Lavrov, 2015; Shtolz and Mishmar, 2023). Interestingly, *atp8* appears to have been lost twice in CBHS: on branches leading to B2 and B5 clades. These losses correlate with losses of the majority of tRNA genes in these lineages and are the only reported losses of *atp8* in Demospongiae (Lavrov et al., 2023).

### Why tRNA genes are retained in mtDNA?

Rapid and recurring losses of tRNA genes in CBHS as well as several other animal groups and many eukaryotes (especially outside of Ophistokonta) raise a question of why at least some tRNA genes have been retained in most mitochondrial genomes. Indeed, a complete replacement of mt-tRNA genes would allow an organism to reduce the size of mtDNA as well as dispose of multiple components involved in mt-tRNA processing, modification, and aminoacylation: changes that should be selectively advantageous. In addition, cy-tRNAs have more conserved structures than their mt counterparts and thus may outperform them in function. Even if selection is too weak to drive tRNA loss, tRNA import, if present, should make mt-tRNA genes redundant and thus subject to accumulation of mutations and, eventually, disappearance. We can think of several answers to this question. First, some mt-tRNAs may have unique functions in mitochondrial translation. Mitochondrial *trnM(cau)* and *trnW(uca)* are prime examples. Because translation within metazoan mitochondria is initiated with N-formylmethionine (fMet), aminoacilated mt-tRNA(Met) should be recognized by mitochondrial methionyl-tRNA formyltransferase (MTFMT) (Bianchetti et al., 1977). Consequently, fMet-tRNAMet is recognized by the mitochondrial translation initiation factor (IF2mt), and together they are recruited to the ribosomal P site to initiate mitochondrial translation (Spencer and Spremulli, 2004). Because fMet is not used in cytosolic translation, cytosolic tRNA(Met) should not be recognized by the MTFMT enzyme.

Similarly, because most animals use a mitochondrial genetic code where UGA codon specifies tryptophan rather than termination of translation, mt-tRNA(Trp) should be able to recognize it along with the standard UGG codon. Again, it would be disastrous to an organism if a cytosolic tRNA(Trp) acquires such function. In fact, *trnW(uca)* and *trnM(cau)* are the only tRNA genes that are retained in the phylum Cnidaria and as well as the demosponge Subclass Keratosa and are among those always retained in the CBHS species. Another explanation, at least for bilaterian animals, is that retention of tRNA genes may be due to the unorthodox structures of encoded tRNAs and their loss of standard recognition features (Kuhle et al., 2020). Indeed, it has been shown that cytosolic and bacterial tRNA synthethases do not recognize mitochondrial tRNAs (Kumazawa et al., 1991; Fender et al., 2012). It is possible that there have been changes in other enzymes that interact with mt-tRNAs, which prevents their replacement by cytosolic counterparts. Third, mt-tRNAs may need to be retained for functions other than carrying amino acids to ribosomes. For example, mt-tRNA(Phe) and mt-tRNA(Val) in mammals are used as a structural RNA within the large subunit of the mitochondrial ribosome, as a replacement for bacterial 5S rRNA (Greber et al., 2014). tRNA genes also act as punctuation marks in the processing of polycistronic transcripts when they are recognized and cleaved from it by processing endonucleases (Rossmanith, 2012). This cleavage not only frees tRNAs but also produces monocistronic transcripts of coding genes that are then utilized in mitochondrial translation (Barchiesi and Vascotto, 2019).

Indeed, the loss of tRNA genes in octocorals disrupts this process and results in bicistronic or tricistronic transcripts (Shimpi et al., 2017). While the functional answers above are possible and even plausible, they are likely incomplete. Indeed, all tRNA genes have been lost in Ctenophora (Pett et al., 2011) and Myxozoa (Takeuchi et al., 2015; **?**), two large groups of non-bilaterian animals, indicating that all tRNA genes are potentially disposable. Even within Bilateria, many (Faure and Casanova, 2006) or nearly all (Barthelemy and Seligmann, 2016) mt-tRNA genes have been lost in chaetognaths. Thus, we may need to look for an alternative explanation. One possibility is that while the loss of mt-tRNA would be beneficial to an organism, tRNA import machinery is simply not available in some groups or some tRNAs can’t be recognized by it. There is an ongoing debate on universality of tRNA import system and on whether the mitochondrial import of tRNA genes occurs in the species without mt-tRNA loss (Alfonzo and Söll, 2009).

Alternatively the loss of tRNAs might be disadvantageos in a long run. As described above, the loss of tRNA genes is often associated with higher rates of sequence evolution and the two animal lineages that have completely lost mt-tRNA genes have the highest rates of mtDNA evolution or have lost mtDNA altogether (Yahalomi et al., 2020).

## Conclusions

Clade B of Haplosclerid Sponges (CBHS) represents a unique system among animals to study the patterns and effects of tRNA loss on mitochondrial genome evolution. While some representatives of this group encode a complete set of tRNA genes required for mitochondrial protein synthesis, others lost as many as 22 tRNA genes. Furthermore, the loss of tRNA genes has occurred independently in several lineages within this group. There has been a large acceleration in the rates of sequence evolution in the lineage leading to B4 and B5 clades as is common in animal mt-genomes missing some tRNA genes. However, other lineages with an extensive loss of tRNAs have not experienced acceleration in rates of mt-sequence evolution. There seems to be no particular relationship between tRNAs encoded in the genome and the codon usage in mitochondrial coding sequences. More mitochondrial and nuclear genomes from this group would be needed to understand the mechanism of tRNA import and driving forces behind mt-tRNA loss.

## Materials and methods

### Dataset construction

Complete mt-genomes of *Niphates digitalis*, *N. erecta*, *Xestospongia muta*, *X. testudinaria* and a nearly complete mt-genome of *Neopetrosia sigmafera* have been assembled and analyzed previously (Wang and Lavrov, 2008; Lavrov et al., 2023). Mitochondrial genome of *Amphimedon queenslandica* was reported by (Erpenbeck et al., 2007).

### New sample acquisition, DNA sequencing and assembly

Samples of *Gelliodes wilsoni* Carballo, Aquilar-Camacho, Knapp & Bell, 2013, *Haliclona (Soestella) caerulea* (Hechtel, 1965), *Haliclona (Gellius) loe* Vicente et al. 2024, *Haliclona (Rhizoniera) pahua* Vicente et al. 2024, *Haliclona (Reniera) kahoe* Vicente et al. 2024 and *Haliclona* sp.14 (H2560) were collected in Kāne’ohe Bay and Ke’ehi Harbor on the island of O’ahu, Hawai’i and preserved in 95% ethanol as described in (Vicente et al., 2021). Samples of two other undescribed *Haliclona* species (TLT723 and TLT785) were collected in California and preserved in 95% ethanol, as described in (Turner and Pankey, 2023). TLT723 was collected from a floating marina dock in Ventura, California on December 14, 2020. Preliminary genetic data from other samples (not shown) indicate that this genotype is found on floating docks throughout Southern California. TLT785 was collected from a tidal pool at Bird Rock, San Diego, California, January 10, 2021. Several samples of this genotype were collected at this location, but this genotype has not been found at any other location to date. Both ”TLT” samples are currently archived in the sponge collation of Thomas Turner at the University of California, Santa Barbara; after additional taxonomic and systemic work, they will be vouchered in a permenant museum collection. Sample of *Haliclona (Haliclona) simulans*(Johnston, 1842) was collected by Christine Morrow from Barhall, Strangford Lough in the east of Northern Ireland on August 9th, 2021 and that of *Haliclona (Halichoclona) plakophila* Vicente, Zea & Hill, 2016 (Sample ID XDPR9) was collected at collected at Old Buoy, La Parguera, Puerto Rico on August 13, 2012. Both were preserved in RNA later solution. Total DNA was extracted from each sample with a standard phenol/chloroform DNA extraction protocol. Whole genome shotgun (WGS) library preparation and sequencing was completed by the DNA Sequencing and Synthesis Facility of the ISU Office of Biotechnology. The WGS libraries were prepared with the Illumina True-Seq PCR free kit. The Hawaiian samples were sequenced on an Illumina HiSEQ 3000 instrument in a 2x100bp paired end run. Samples from California were sequenced on an Illumina NovaSeq6000 instrument using a SP flow cell in a 2x150bp paired end run.

DNAseq data was assembled using the SPAdes 3.14.0 (Prjibelski et al., 2020). In some cases a better assembly was achieved using subsamples of data with seqtk-1.4 (r122) program (https://github.com/lh3/seqtk). Contigs containing mitochondrial sequences were recognized by their sequence similarity to mtDNA of known haplosclerid sponges with fasta-36.3.8 (Pearson and Lipman, 1988). Potential problems in the assemblies were checked by mapping raw reads to the assembly with BOWtie v.1.2 (Langmead et al., 2009).

### Mt-genome annotation

Mt-genomes were annotated using mfannot online server https://megasun.bch.umontreal.ca/apps/mfannot/ (Lang et al., 2023). Additional tRNA genes were searched with the tRNAscan-SE v.2.0.6 (Chan et al., 2021) using Organellar Search Mode (-O) option (invoking maximum sensitivity mode) and, separately, using the -C (cove) option. *trnI(cau)*… *trnS(gcu)* was mis-identified in *Gelliodes wilsoni* as the second trnT(ugu) and its identity was corrected manually based on sequence similarity with closely related species. rRNA boundaries were checked and adjusted manually based on interspecific alignment and RNA-seq data (when available).

### Phylogenetic analysis

#### tRNA genes

mt-tRNA sequences predicted by tRNscan-SE were aligned with the cmalign program from the Infernal 1.1.3 package (Nawrocki and Eddy, 2013) using -g –notrunc –outformat AFA options and the TRNAinf-1415.cm covariance model. Neighbor joining phylogenetic analysis of all mt-encoded tRNAs was conducted in PAUP Version 4.0a (build 168) (Swofford, 2002). p-distances were used to avoid overfitting (Sullivan and Swofford, 2001).

#### Coding sequences

Coding sequences from 13 mitochondrial protein genes were aligned with Mafft v7.475 (Katoh and Standley, 2013) using L-INS-i strategy. Conserved blocks within the alignments were selected with Gblocks 0.91b (Talavera and Castresana, 2007) using relaxed parameters (parameters 1 and 2 = 0.5, parameter 3 = 8, parameter 4 = 5, all gap positions in parameter 5). Cleaned alignments were concatenated into a supermatrix containing 3,696 amino acid positions for 24 species. Four chains in Phylobayes-MPI (Lartillot et al., 2013) were run for *>*12,000 cycles until the mean standard deviation of bipartition frequencies was *<*0.1.

#### Mapping gene loss

We used the Dollo parsimony principle, which assumes the irreversible loss of characters, to manually map gene loss on the phylogenetic tree. The results were adjusted based on the analysis of tRNA gene phylogenetic relationships, which revealed changes in some anticodon identities.

### Mt-sequence evolution

We used Composition profiler (Vacic et al., 2007) to identify significant changes in amino acid frequencies in comparison to *Xestospongia muta*. We used within-group correspondence analysis of codon usage data (Charif et al., 2004) to investigate changes in synonymous codon usage. The published R script (https://pbil.univ-lyon1.fr/datasets/charif04/) has been slightly modified to use local files and a minimally modified genetic code (NCBI genetic code 4)

### Identification of *atp8* and *atp9* in the nuclear genomes

We used mmseqs2 (Steinegger and Söding, 2017) to identify contigs with similarity to *Xestospongia muta atp8* and *atp9* in the transcriptome and draft genome sequence from *Niphates digitalis* (unpublished data). In order to determine if the location of *atp9* in the *Niphates digitalis* genome was the same as in *Amphimedon queenslandica*, contigs containing *atp9* were extracted from the *N. digitalis* and from the *A. queenslandica* nuclear genome assemblies. Dot plots were made comparing the contigs using YASS: genomic similarity search tool (Noe and Kucherov, 2005).

## Acknowledgments

This work was supported by the Division of Environmental Biology at the National Science Foundation [DEB No. 0829783 to D.V.L.] and by the Presidential

Interdisciplinary Research Seed (PIRS) Grant from Iowa State University to [D.V.L.]. Postdoctoral work of J.V. was supported by the Division Of Ocean Sciences at the National Science Foundation [OCE No. 2048457]. We thank Christine Morrow for collecting a sample of *Haliclona simulans* and Ehsan Kayal for helpful comments on an earlier version of this article.

## Data Availability

The WGS read files can be found under the NCBI BioProject PRJNA1075689; assembled and annotated mitochondrial genomes have been deposited under accession numbers XXXXXXXX-YYYYYYYY.

